# Buildings act as barriers to insects, decoupling the bacterial-dominated and insect-dominated phases of carcass decomposition

**DOI:** 10.64898/2026.11.25.769181

**Authors:** Isabelle Perron, Lena Lutz, Gaétan Moreau

## Abstract

The dynamics of necrophagous insect communities and the decomposition of the cadavers they colonise can be modulated by ecological filtering limiting insect access. We examine how different degrees of confinement influence the decomposition of animal carcasses, their colonisation by necrophagous insects, and the link between these two processes. To this end, vertebrate carcasses were placed either outdoors or in sheds with open or closed windows in a replicated experiment. Over two weeks, decomposition and insect abundance were documented daily. The results indicate that increasing confinement altered insect-mediated decomposition processes, delaying the insect-dominated phase while leaving the bacterial-dominated phase unaffected. Thus, in indoor environments, a dissociation of the decomposition sequence normally observed outdoors was observed due to an increasing delay in the colonisation of most insects with increasing confinement. This indicates that buildings can act as ecological filters that structure the accessibility of carcasses to necrophagous insects. In addition, these findings suggest that cadaver decomposition in indoor environments follows a distinct trajectory, with direct consequences for postmortem interval estimations.

## Introduction

Constraints to colonisation, such as geographic isolation, microclimatic conditions or mechanical barriers, are major determinants of ecological succession as they modulate species establishment and the trajectory of succession (Valiela 1974; Hughes & Fahey 1988; Turner et al. 1998; Yan et al. 2020). Although plant succession is often used as the reference model, this conceptual framework of ecological succession also applies to the succession of heterotrophic necrophagous organisms observed during large vertebrate carcass or human cadaver decomposition (Michaud et al. 2015). In this context, constraints on the colonisation of decomposer organisms may also affect the rate of decomposition of necromass (Payne 1965; Simmons et al. 2010; Reibe & Madea 2010; Charabidze et al. 2015), although the mechanisms by which these factors modulate decomposition dynamics remain largely unresolved.

Buildings are among the principal barriers to the colonisation of cadavers by necrophagous insects, restricting access to remains and altering microclimatic conditions, decomposition processes, and insect physiology and development. In Canada and elsewhere, most homicides occur in enclosed environments where colonisation constraints vary according to the degree of confinement imposed by buildings (Catanese & Bloom 2002; Pohjoismäki et al. 2010; Syamsa et al. 2017; Lutz et al. 2018; Statistics Canada 2022). Despite this, the effects of buildings remain poorly documented, because forensic entomology research has largely been conducted outdoors. The few studies conducted indoors (e.g., Reibe & Madea 2010; Anderson 2011; Aly et al. 2017; Almutawa et al. 2024) have observed delayed insect colonisation and oviposition, particularly by dipterans, as well as delayed decomposition in carcasses placed inside buildings compared with those placed outdoors. However, these observations do not permit causal inference because the studies are affected by pseudoreplication (Michaud et al. 2012; Moreau et al. 2025).

Carcass or cadaver decomposition generally begins with a putrefaction phase initiated by endogenous bacteria (Janaway et al. 2009). This bacterial-dominated phase is followed by a insect-dominated phase during which blowfly larvae (Diptera: Calliphoridae), as well as other dipterans and beetles (Coleoptera), become the main agents of necromass degradation (Mégnin 1887; Campobasso et al. 2001). These phases differ primarily in the relative contribution of bacteria and insects to decomposition. Although decomposition is traditionally described using qualitative stages, the TBS method (Megyesi et al. 2005; Keough et al. 2017) provides a finer and more continuous measure of decomposition progression (Simmons et al. 2010; Moffatt et al. 2016; Lutz et al. 2019), allowing decomposition rates and their temporal variation to be related directly to continuous measures of insect colonisation.

Herein, we examined how different degrees of confinement imposed by human buildings influence carcass decomposition and colonisation by necrophagous insects. We test the hypothesis that vertebrate decomposition is influenced by constraints on colonisation, acting as ecological filters that modulate the succession dynamics of necrophagous insects and the rate of entomological decomposition. The conceptual framework of assembly filters (Weiher & Keddy 1995) postulates that local communities result from successive filters that restrict the local pool of species. This may also slow colonisation by modulating the probability and rate of species establishment. From this hypothesis arise three predictions, namely that (P1) increasing degrees of confinement of buildings surrounding carcasses reduce or delay access by certain necrophagous insect taxa, (P2) the presence and abundance of certain necrophagous taxa are closely associated with the rate of decomposition quantified by the TBS method, and (P3) increasing confinement slows the insect-dominated decomposition phase without altering the bacterial-dominated decomposition phase. Thus, under outdoor conditions, decomposition should follow a relatively continuous progression from autolysis and putrefaction to insect-mediated processes. In contrast, in confined environments, decomposition should be structured into two distinct phases: an initial phase marked by a delay in insect arrival, followed by a pronounced acceleration of decomposition once colonisation is established. This study also allowed us to test prediction (P4) that the contribution of insects to decomposition rates is not constant over time, but increases progressively as entomological colonisation becomes established. Although intuitive, this prediction has not previously been tested experimentally.

## Materials and Methods

### Experimental Design

The study was conducted in an abandoned 2-ha agricultural field located in Cocagne, New Brunswick, Canada (46° 20.364’ N, 64° 39.724’ W). A detailed description of the predominantly herbaceous plant community can be found in LeBlanc et al. (2021). During this study, climatic conditions were characterized by mean daytime and nighttime temperatures of 27.2 °C and 16.1 °C, respectively, as well as dry conditions with cumulative precipitation of 28.7 mm. Temperature and humidity inside the sheds and around outdoor cages are described in Dixon et al. (2026).

Ten domestic pigs (*Sus scrofa domesticus*) weighing 21.32 ± 2.18 kg were obtained from a certified pig farm located 60 km from the study site. They were euthanized using a captive bolt gun placed on their forehead. Immediately afterward, each carcass was placed in two thick airtight plastic bags to prevent insect colonisation during transport to the study site. Upon arrival, carcasses were placed in experimental units at least 40 m apart (Fig. A1) to reduce cross-contamination and competition among insects for carcasses (Matuszewski et al. 2020).

Eight of the experimental units consisted of metal sheds measuring 1.83 m × 1.22 m, Shed-in-a-Box™ brand, model no. SBS64 by Arrow Storage Products. Each shed was placed on a rectangular vinyl floor extending 10 cm beyond the perimeter of the shed to prevent the formation of gaps between the structure base and the floor. The doors were sealed with adhesive tape, which was regularly replaced due to daily opening for sampling. To prevent intrusion by large vertebrate scavengers, cinder blocks were used to keep the doors closed between sampling events. To allow air circulation, while limiting insect entry, 10 × 25 cm vents were cut into the upper part of the door and at the bottom of the opposite wall, and were covered with fine tulle held in place with glue and adhesive tape. The sheds were insulated with a thick layer of spray foam, thereby restricting insect access and delaying colonisation. A rectangle of white tarp attached with adhesive tape replaced a 39 × 58 cm panel on the shed, creating a closed window. For sheds with an open window, a 2-cm-high × 58-cm-wide slit was left at the bottom of the rectangle. A pig carcass was placed in each shed in the corner opposite the door and oriented along a slope of approximately 2% to prevent decomposition fluids from flowing toward the entrance. The sampling area was the whole cabin.

Two other experimental units were delimited by a geotextile membrane measuring 70 × 140 cm placed on the ground and covered with 2 cm of vermiculite to allow pupation of certain insects. A carcass was placed at the center of the membrane and covered with a 122 × 61 × 61-cm wooden frame covered with galvanized steel mesh wire with 3-cm regular hexagonal openings. Cages were held in place with elastic cords and cinder blocks to prevent vertebrate scavenging. The membrane allowed drainage of carcass fluids and precipitation, delimited the sampling area, and facilitated the collection of insects that burrow into the soil.

The control treatment was replicated twice, whereas the shed treatments were replicated four times each for open and closed windows. This decision was based on previous data (e.g., Michaud & Moreau 2013, 2017; LeBlanc et al. 2021; Losier et al. 2023) showing low intra-treatment variability for control carcasses placed outdoors in this study area. We expected that shed treatments, which had not previously been investigated, would exhibit higher variability in colonisation, justifying greater replication to ensure adequate statistical power (Krebs 1999).

### Sampling

Insect communities and carcass decomposition were documented and sampled ∼3–4 h postmortem, then daily for the following 14 days. Most carcasses had reached the dry stage by day 14. The order of carcass visitation was randomized daily.

During shed sampling, one person remained outside to take notes and close the door, while another person, equipped with a headlamp, entered the shed through the lower part of the door and conducted observations and collected samples. When the door was opened, the person outside also recorded the number of escaping blowflies, which consisted of only a few individuals throughout the study; no insects were observed entering through the door during the experiment. For cage sampling, cages were approached slowly and carefully so as not to disturb the insects. A count of insects was then performed before carefully lifting the cage. After waiting for one to two minutes for insects to resume their activity, samples were collected and observations completed.

Entomological observations included recording the presence of fly larvae and counting individuals belonging to each recognizable taxonomic group within the sampling area, following the method of Losier et al. (2023). Most beetle specimens were identified to species directly in the field. Taxa that could not be identified in the field, such as Calliphoridae, small dipterans, and small beetles, were collected with an entomological net or forceps and preserved in 70% ethanol. To minimize disturbance to the insect community, no more than 10% of the observed individuals within a given taxon were collected. Photographs and videos of the entire carcass, limbs, the torso, and the head were also taken.

In the laboratory, photographs and videos were analyzed following the method of Keough et al. (2017) to obtain the TBS for each carcass at each sampling event. In addition, sampled specimens were identified using taxonomic keys (e.g., McAlpine 1981; Jones et al. 2019) and the reference collection housed at the Université de Moncton. Four insects that could not be reliably identified based on morphology alone were sent to the Institute of Legal Medicine in Frankfurt, Germany, for DNA barcoding. Eight necrophagous insect taxa counted more than 50 times were selected for further analysis. Among Diptera, these include adult Calliphoridae, Calliphoridae larvae, adult Muscidae, and small adult Diptera, primarily Piophilidae and Sepsidae. Among Coleoptera, these include adult *Creophilus maxillosus*, adult *Necrodes surinamensis*, larvae of *Necrodes surinamensis*, and adult *Oiceoptoma noveboracense*.

### Statistical Analyses

All analyses were conducted in R 4.5.2 (R Core Team 2025). The rate of change in TBS was calculated for each carcass by computing the difference in total TBS between two successive sampling events. To highlight the relationship among postmortem interval (PMI), decomposition, insect taxa, and experimental treatments, we fitted a series of generalized additive models (GAMs) using the *mgcv* package. Each variable was modeled as a function of PMI using a treatment-specific smoothing term (s(PMI, by = treatment)) and the random effect of experimental unit (s(unit, bs = “re”)). Models were fitted with Poisson, binomial, or negative binomial distributions depending on the distribution of the response variable. For each model, predictions and their confidence intervals were generated for each treatment. In parallel, cross-correlation functions (CCF) were calculated between the rate of change in TBS and each insect taxon to assess temporal lag between insect activity and decomposition progression.

Daily abundances of the eight taxa showing significant CCFs (see Results) were transformed using the Hellinger method to standardize abundance data and incorporate larval presence while reducing differences in scale among variables. A principal component analysis was performed on the transformed data using the *vegan* package to obtain a first canonical axis (PC1_insects), which explained 58% of the variation in insect composition and abundance, was retained as a synthetic measure of insect colonisation intensity. This variable was then included in a model of the rate of change in TBS including a smoothed effect of postmortem interval (s(PMI)), a random effect for each carcass (s(unit, bs = “re”)), an insect effect varying with time (s(PMI, by = PC1_insects)), a treatment effect varying with time (s(PMI, by = treatment)), and a tensor-product interaction among PMI, insect colonisation intensity, and treatments (ti(PMI, PC1_insects, by = treatment)). Because the rate of change in TBS included zero values, a Tweedie family with a log link was used. All additive models were fitted by REML.

## Results

### Dynamics of decomposition and insect activity

Across most taxa, decomposition and insect dynamics showed pronounced non-linear responses to PMI, and these responses differed consistently among outdoor, open-shed, and closed-shed treatments (Table 1; Figure 1a–i). For all variables analyzed, except the abundance of *O. noveboracense*, the treatment-specific smoothing functions were significant (Table 1), and their effective degrees of freedom (*edf*) were greater than 2. This indicates that the effect of PMI differed among treatments and that the relationship between PMI and the response variable was non-linear. In the case of *O. noveboracense*, the smoothing functions associated with outdoor conditions and closed sheds were significant, indicating non-linear temporal dynamics. However, the smoothing function for open sheds was not significant, suggesting that for this treatment, PMI did not explain a detectable proportion of the variation in species abundance.

**Figure 1.**
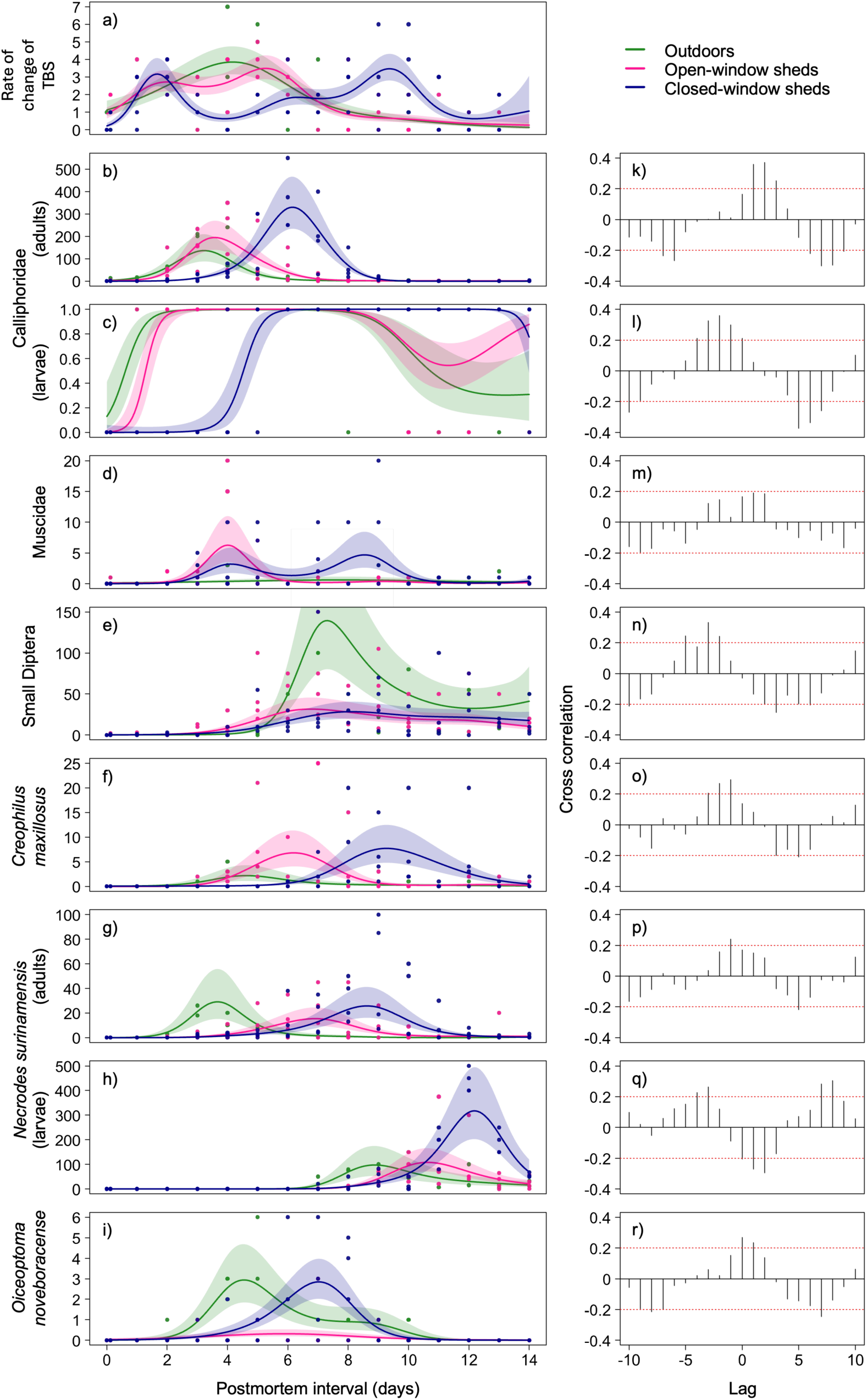
Generalized additive models of a) the rate of change in decomposition, b) adult abundance of Calliphoridae, c) larval presence of Calliphoridae, d) Muscidae abundance, e) small Diptera abundance, f) *Creophilus maxillosus* abundance, g) adult abundance of *Necrodes surinamensis*, h) larval abundance of *Necrodes surinamensis*, and i) *Oiceoptoma noveboracense* abundance as a function of postmortem interval for each treatment, namely outdoors, in sheds with an open window, and in sheds with a closed window. Shaded areas around the fitted curve represent standard errors. Figures k-r show cross-correlations between the abundance or presence of each insect taxon shown in panel b-i and the rate of change in decomposition shown in panel a.

**Table 1.** Data distribution, model r² and chi-square values for the smoothed effect of the treatment × postmortem interval (PMI) interaction for each variable analyzed using generalized additive models. The models were specified as: Variable = s(PMI by treatment) + s(experimental unit). Only taxa observed more than 50 times were analysed.

| Variable | Distribution | *r*<sup>2</sup> | Outdoors × PMI | Open-window cabins × PMI | Closed-window cabins × PMI |
|---|---|---|---|---|---|
| Rate of TBS change | Poisson | 0.41 | 34.60** | 48.18** | 39.23** |
| Diptera |  |  |  |  |  |
| Calliphoridae (adults) | Negative binomial | 0.66 | 94.40** | 250.21** | 273.43** |
| Calliphoridae (larvae) | Binomial | 0.72 | 9.30* | 9.53* | 9.60* |
| Muscidae | Poisson | 0.44 | 7.83* | 94.81** | 65.48** |
| Small Diptera | Negative binomial | 0.46 | 52.53** | 89.87** | 82.56** |
| Coleoptera |  |  |  |  |  |
| *Creophilus maxillosus* | Negative binomial | 0.31 | 10.60* | 37.12** | 39.79** |
| *Necrodes surinamensis* (adults) | Negative binomial | 0.64 | 34.52** | 47.75** | 69.24** |
| *Necrodes surinamensis* (larvae) | Negative binomial | 0.68 | 38.32** | 71.00** | 102.57** |
| *Oiceoptoma noveboracense* | Negative binomial | 0.46 | 36.07** | 6.50 | 47.13** |

\* 0.01 < *P* < 0.05
\*\* *P* < 0.01

Increasing carcass confinement generally delayed insect activity peaks and altered taxon-specific abundance patterns (Fig. 1a-i). The smoothing functions for the rate of change in TBS increased and decreased relatively uniformly in the outdoor treatment (Fig. 1a). In open sheds, the overall pattern remained similar but showed a tendency to split into two peaks. This phenomenon was even more pronounced in closed sheds, where at least two distinct and well-separated peaks were visible. For Calliphoridae adults, abundance increased and peak occurrence was progressively delayed with increasing carcass confinement (Fig. 1b). The presence of Calliphoridae larvae occurred later as the level of confinement increased (Fig. 1c). Muscidae were rarely observed on outdoor carcasses. However, they were very abundant on carcasses in sheds, with an early abundance peak in open sheds and a later peak in closed sheds (Fig. 1d). Small dipterans did not exhibit a temporal shift among treatments, but their abundance was higher in the outdoor treatment (Fig. 1e). *Creophilus maxillosus* showed an abundance pattern very similar to that of adult Calliphoridae (Fig. 1f). Adult *N. surinamensis* also exhibited an increasing delay in abundance with increasing confinement, although their abundance was slightly lower in the open shed treatment (Fig. 1g). Larvae of *N. surinamensis* followed the same temporal delay, but increased dramatically in abundance in closed sheds (Fig. 1h). Finally, *O. noveboracense* was almost absent from the open shed treatment and showed later colonisation in closed sheds than on carcasses exposed outdoors (Fig. 1i).

### Association between insects and decomposition

Cross-correlation analyses revealed that, for all taxa except Muscidae, the relationship between insect abundance or presence and the rate of change in decomposition varied significantly across temporal lags, with correlations alternating between positive and negative values (Fig. 1k-r). All series were stationary. Several taxa showed positive correlations at negative lags (Figs. 1l, n, q) or positive correlations at positive lags (Fig. 1k), indicating a dynamic and phase-shifted relationship with the decomposition process. Three taxa showed a positive association at lag 0 (Figs. 1o, p, r), while one taxon deviated from this general pattern by showing a negative correlation at lag 0 (Fig. 1q), suggesting an immediate inverse association with the rate of change in decomposition.

An additive model was developed to assess whether the contribution of insects to the rate of decomposition varied over time and across treatments. This model explained a larger proportion of the variability in the rate of change in decomposition (47%) than the previous model, which did not include insect-related variables (Table 2). The term s(PMI) indicated that the rate of change in decomposition depended strongly on postmortem interval. The term s(PMI):PC1_insects revealed that entomological activity played a significant role in shaping decomposition dynamics. In addition, the effect s(PMI):outdoor showed that this relationship differed in the outdoor treatment. Finally, a marked interaction among PMI, insect activity, and the closed-shed treatment was detected (ti(PMI, PC1_insects):closed shed), whereas no significant interaction was observed for the outdoor or open-shed treatments.

**Table 2.**
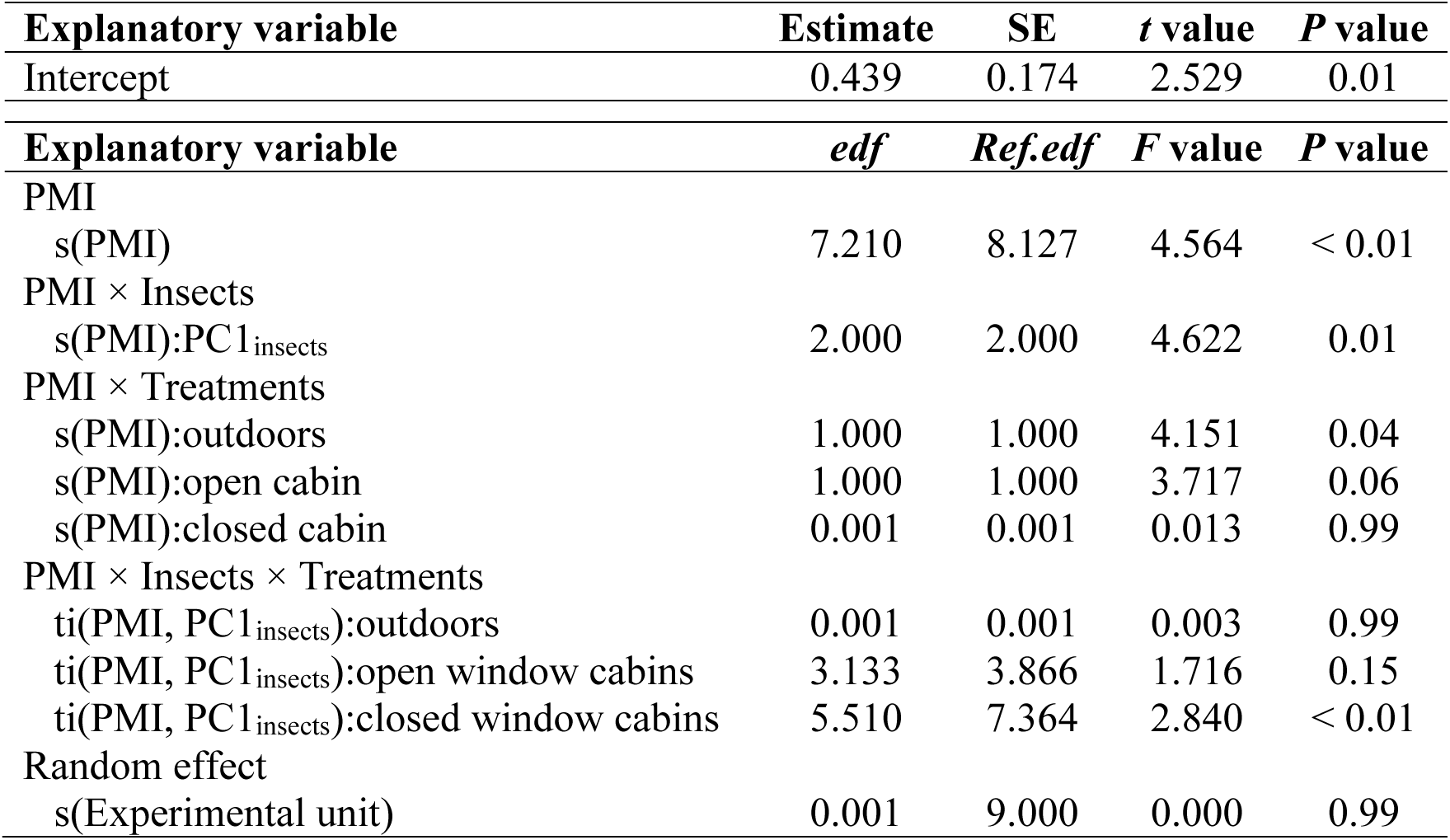
ANOVA table for regressions of the generalized additive mixed model (GAMM) predicting decomposition rate by TBS as a function of insects and PMI.

## Discussion

This study demonstrates that carcass confinement alters necrophagous insect succession, delays the insect-dominated phase of decomposition, and modifies the relationship between insect activity and decomposition dynamics. Specifically, we show that (P1) increasing degrees of confinement of the structures surrounding carcasses reduce or delay access by certain necrophagous insect taxa, (P2) the presence and abundance of certain necrophagous insects are closely associated with decomposition rates, and increasing confinement slows the insect-dominated decomposition phase without altering the bacterial-dominated decomposition phase. (P3) increasing confinement slows the insect-dominated decomposition phase without altering the bacterial-dominated decomposition phase. While decomposition was not prevented, delayed colonisation by key necrophagous taxa modified the timing of decomposition and postponed the onset of the insect-dominated decomposition phase. This provides support for our hypothesis that cadaver decomposition is influenced by constraints on colonisation, acting as ecological filters that modulate the succession dynamics of necrophagous insects and entomological decomposition. This study also validates the prediction (P4) that the contribution of insects to decomposition rates increases progressively as entomological colonisation becomes established.

Our study indicates that in outdoor environments, the progression of decomposition is continuous, whereas in indoor environments, decomposition becomes gradually more discontinuous with increasing levels of confinement. Two distinct decomposition modes emerge: a first that corresponds to the bacterial-dominated phase, and a second that we attribute to the insect-dominated phase (Fig. 2). Although the two phases may be influenced by abiotic conditions (Carter et al. 2008; Voss et al. 2008), it is unlikely that decomposition was slowed by the microclimatic conditions of the sheds. On the contrary, the slightly warmer (≈ +5 °C) and more humid (5–10%) conditions observed in the sheds during the entomological phase (Dixon et al. 2026) would be expected to favor bacterial activity and insect feeding. Thus, the decoupling observed between the two phases is most consistent with the documented delay in entomological colonisation, rather than with climatic inhibition of decomposition processes.

**Figure 2.**
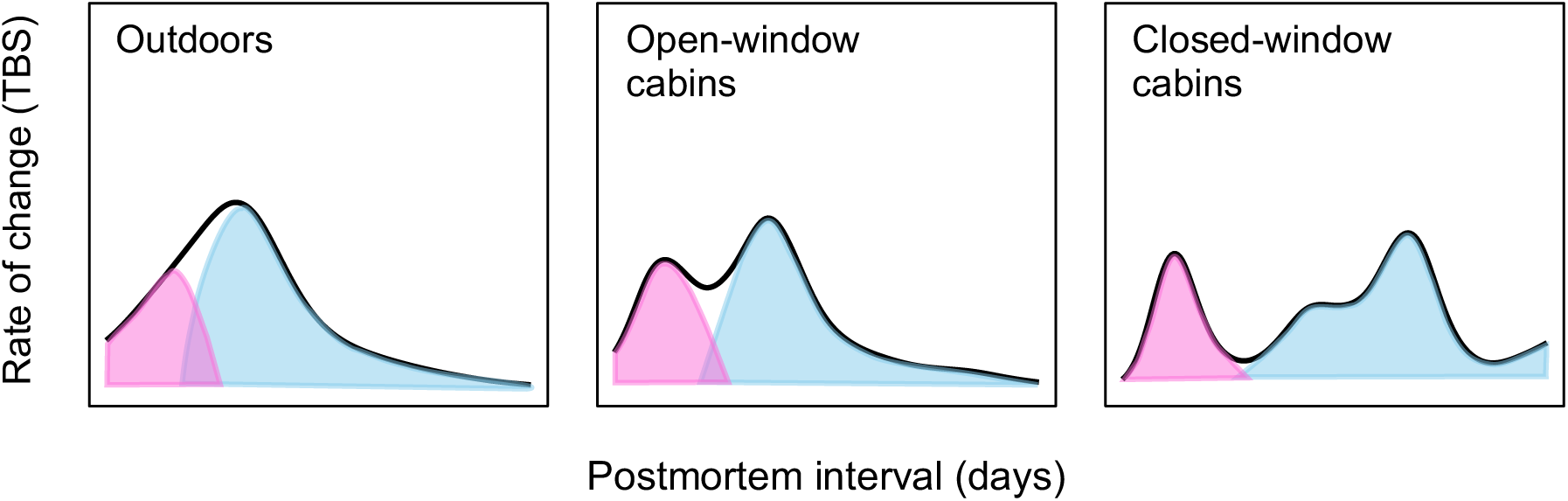
Theoretical representation of the bacterial-dominated phase (pink) and insect-dominated phase (blue) for carcasses placed a) outdoors, b) in sheds with an open window, and c) in sheds with a closed window.

Although refinements to the TBS method have been proposed for indoor human decomposition, particularly to account for insect infestations and extensive desiccation (Ceciliason et al. 2018), these limitations are unlikely to have influenced our results because the divergence in decomposition trajectories and colonisation delays we documented occurred before either process became prominent. We therefore retained the approach of Keough et al. (2017), which is better suited to porcine models.

### Confinement as a taxon-specific filter of insect colonisation

The results also highlighted the important effect of buildings on the ability of Diptera and Coleoptera to colonise carcasses. Calliphoridae, which are pioneer colonisers whose larvae often constitute the main driver of entomological decomposition (Payne 1965; Anderson & Cervenka 2001), took longer to access more confined environments. Paradoxically, their maximum abundance was higher in closed sheds, possibly due to their inability to leave once they had entered. The delay in Calliphoridae larval colonisation contributed to the observed deceleration of entomological decomposition (Campobasso et al. 2001; Thümmel et al. 2023).

Muscidae, an opportunistic taxon attracted to a wide diversity of substrates (Skidmore 1985) and a presumably weaker competitor than obligate necrophages, were underrepresented in our samples and showed only a limited association with decomposition. This taxon colonised open sheds earlier than closed sheds, possibly due to the low level of inter-dipteran competition in early decomposition. Small dipterans, mainly Sepsidae and Piophilidae, were not delayed by the sheds, but exhibited lower abundances indoors in comparison to outdoor conditions. Confinement likely did not delay their initial arrival because these species are highly mobile and capable of rapidly entering enclosed environments through very small openings (Freidberg 1981). However, because confinement reduces available space, competition with dominant dipterans (e.g., Calliphoridae) may be exacerbated in sheds.

*Creophilus maxillosus* and both larval and adult stages of *N. surinamensis*, all of which prey on Calliphoridae larvae, also exhibited delayed colonisation with increasing confinement. This delay can be explained by their difficulty in penetrating indoor environments due to their larger size (12 to 25 mm), as well as by the delayed colonisation of their primary food source, dipteran larvae. *Oiceoptoma noveboracense*, another predatory and necrophagous beetle, was also delayed, but only in closed-window sheds, possibly for reasons similar to those affecting the previous taxa.

The eventual colonisation of insects in closed-window sheds does not result from accidental introductions associated with handling, as opening the door resulted instead in a negligible escape of flies from the sheds. As mentioned previously, tiny insects can exploit microcracks in the structure of building that are invisible to the naked eye (Canadian Conservation Institute 2019). This could make it easier for them to access carcasses within buildings (Guo et al. 2026). Calliphoridae and other large dipterans are also capable of exploiting holes in insulation. As for beetles, they possess strong mandibles and elytra that could potentially allow them to penetrate certain insulating materials.

### Association between insects and decomposition

Most previous studies have examined insect community dynamics relative to decomposition stages or TBS, whereas the present study focuses on the rate of change of decomposition itself, providing a process-oriented view of carrion decomposition dynamics. Using cross-correlations, we have shown that most taxa track decomposition through temporally lagged responses. The lagged positive correlations suggest that certain groups, including small dipterans, Calliphoridae larvae, and adult Calliphoridae are temporally associated with previous changes in decomposition rate occurring several days earlier rather than to its instantaneous rate of change. Adult Staphylinidae showed a direct and positive association with the rate of change in decomposition, indicating close synchronization with its instantaneous rate, likely linked to their direct dependence on the presence of other necrophagous insects on which they feed (Tantawi et al. 1996). In contrast, larvae of *N. surinamensis* stood out by showing a direct negative association with the rate of decomposition change. This atypical pattern is consistent with their ecology, as opportunistic consumers capable of feeding on both animal tissue and decomposers (Ratcliffe 1972), their local abundance may increase at times when decomposition slows, for example when exploitable biomass decreases or when competing resources become scarce. Conversely, this pattern could alternatively reflect an effect of maggot predation on decomposition rates, although the present study does not allow this mechanism to be tested directly. Overall, these results highlight that trophic position and mode of resource exploitation influence how different taxa are temporally linked to the decomposition process.

Our results also show that the influence of the insect community on the progression of decomposition depends on the experimental treatment. This influence was relatively stable over time outdoors and in open sheds. In contrast, in closed sheds, the interaction between PMI and the insect community intensity was significant, indicating that the relationship between entomological activity and decomposition changes over time. This suggests that the closed-shed treatment profoundly alters decomposition dynamics, likely by limiting insect access and slowing entomological activity. Furthermore, these analyses reveal that insects play a major role in the progression of the second part of the decomposition process, but this role is strongly modulated by confinement conditions. Nevertheless, these results should be extrapolated with caution, as the sheds are not representative of all enclosed structures, such as houses, suitcases, barrels, or cars. The purpose of the sheds was not to recreate these situations, but rather to create a controlled environment with different degrees of confinement to investigate the study of the ecological mechanisms occurring under those conditions.

### Forensic implications

This study, based on a replicated experimental design, demonstrates that increasing confinement leads to a growing temporal dissociation between the bacterial-dominated and insect-dominated phases of decomposition and challenges the direct extrapolation of dynamics observed outdoors to confined environments. By highlighting the structuring effect of confinement on the dynamics of entomological colonisation and on the temporal contribution of insects to decomposition, this study may contribute to a better interpretation of homicide cases involving cadavers found indoors and may provide a mechanistic framework to improve the estimation of the postmortem interval. It thus underscores the necessity to integrate the degree of confinement of death scenes into forensic entomological analyses.

## Acknowledgments

The authors would like to thank Cara Dixon, Denis R. Boudreau, Mélodi Lagacé, Adriana García del Ángel, Gabriel Mazerolle, Jacob Moreau, and Jean-Pierre Privé for help with field work. This research was supported by the Natural Sciences and Engineering Research Council of Canada (NSERC) and MITACs.

## Ethics approval

The pigs used in this study were euthanized in accordance with Canadian animal welfare regulations and the guidelines of the Canadian Council on Animal Care (CCAC) for livestock intended for human consumption. According to Université de Moncton guidelines, animal ethics approval was not required because this study did not involve live animals.

## Appendix

**Figure A1.**
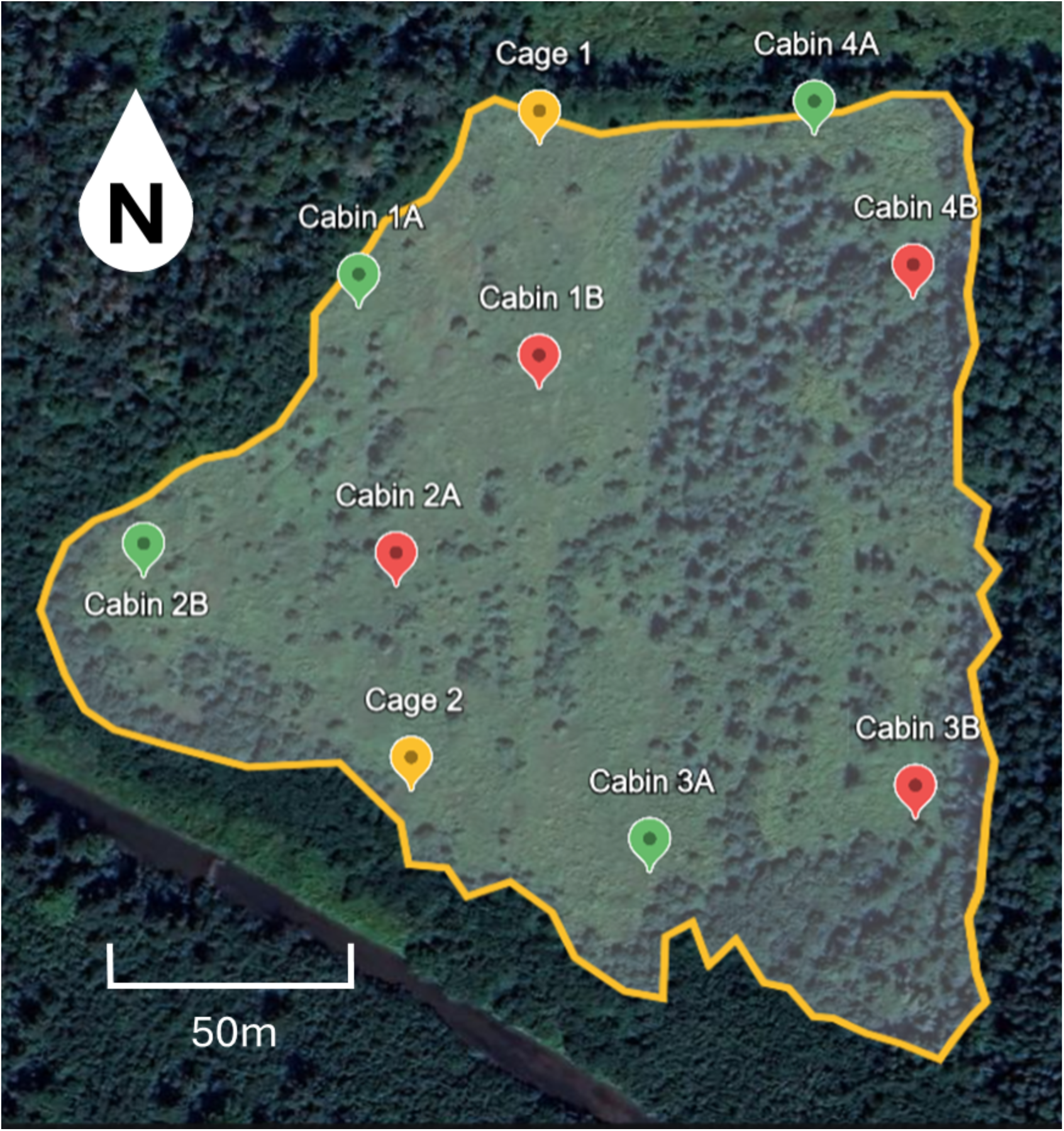
Map of the study site in Cocagne, New Brunswick, Canada (46° 20.364′ N, 64° 39.724′ W), showing carcasses placed outdoors (yellow), carcasses located in sheds with open windows (green), and carcasses located in sheds with closed windows (red).

## References

Almutawa, M. Y., Al-Khalifa, M. S., Al-Dhafer, H. M., Abdel-Dayem, M. S., Ebaid, H., & Ahmed, A. M. (2024). Forensic investigation of carcass decomposition and dipteran fly composition over the summer and winter: A comparative analysis of indoor versus outdoor at a multi-story building. Journal of Medical Entomology, 61(4), 877–890. 10.1093/jme/tjae039

Aly, M. Z. Y., Osman, K. S. M., Galal, F. H., & Ali, G. H. M. (2017). Comparative study on outdoor and indoor forensic insects encountered on rabbit corpses in upper Egypt. IOSR Journal of Pharmacy and Biological Sciences, 12(03), 41–54. 10.9790/3008-1203074154

Anderson, G. S., & Cervenka, V. J. (2001). Insects associated with the body: their use and analyses. Advances in forensic taphonomy (pp. 173–200). CRC Press.

Anderson, G. S. (2011). Comparison of decomposition rates and faunal colonization of carrion in indoor and outdoor environments. Journal of Forensic Sciences, 56(1), 136–142. 10.1111/j.1556-4029.2010.01539.x

Canadian Conservation Institute. (2019). Detecting infestations: Facility inspection procedure and checklist. *Government of Canada*. https://www.canada.ca/en/conservation-institute/services/conservation-preservation-publications/canadian-conservation-institute-notes/detecting-infestations.html

Campobasso, C. P., Di Vella, G., & Introna, F. (2001). Factors affecting decomposition and Diptera colonization. Forensic Science International, 120(1-2), 18–27.

Carter, D. O., Yellowlees, D., & Tibbett, M. (2008). Temperature affects microbial decomposition of cadavers (Rattus rattus) in contrasting soils. Applied Soil Ecology, 40(1), 129–137. 10.1016/j.apsoil.2008.03.010

Catanese, G., & Bloom, T. (2002). Recovery of a mummified pregnant woman from a 55-gallon drum more than 30 years after her death. The American Journal of Forensic Medicine and Pathology, 23(3), 245–247.

Ceciliason, A.-S., Andersson, M. G., Lindström, A., & Sandler, H. (2018). Quantifying human decomposition in an indoor setting and implications for postmortem interval estimation. Forensic Science International, 283, 180–189. 10.1016/j.forsciint.2017.12.026

Charabidze, D., Hedouin, V., & Gosset, D. (2015). An experimental investigation into the colonization of concealed cadavers by necrophagous blowflies. Journal of Insect Science, 15(1), 149. 10.1093/jisesa/iev129

Dixon, C., Perron, I., Lutz, L., Boudreau, D. R., Moreau, G. (2026). A window of opportunity: decomposition and insect colonization of carcasses under outdoor and enclosed conditions. *Under review*.

Freidberg, A. (1981). Taxonomy, natural history and immature stages of the bone-skipper, Centrophlebomyia furcata (Fabricius) (Diptera: Piophilidae, Thyreophorina). Insect Systematics & Evolution, 12(3), 320–326. 10.1163/187631281794709728

Guo, Y., Luo, Y., Ma, Y., Luciano, A., Huang, J., Li, Y., Wang, S., Niu, Y., Li, Z., Cai, J., & Meng, F. (2026). Wings on concealed corpse: The forensic importance of scuttle flies (Diptera: Phoridae). Frontiers in Insect Science, 5, 1727384. 10.3389/finsc.2025.1727384

Hughes, J. W., & Fahey, T. J. (1988). Seed dispersal and colonization in a disturbed northern hardwood forest. Bulletin of the Torrey Botanical Club, 115(2), 89–99. 10.2307/2996139

Janaway, R. C., Percival, S. L., & Wilson, A. S. (2009). Decomposition of human remains. Dans: Microbiology and Aging: Clinical Manifestations, pp. 313–334. Totowa, NJ: Humana Press.

Jones, N., Whitworth, T., & Marshall, S. A. (2019). Blow flies of North America: Keys to the subfamilies and genera of Calliphoridae, and to the species of the subfamilies Calliphorinae, Luciliinae and Chrysomyinae. Canadian Journal of Arthropod Identification. 39: 191 pp. doi:10.3752/cjai.2019.39

Keough, N., Myburgh, J., & Steyn, M. (2017). Scoring of decomposition: a proposed amendment to the method when using a pig model for human studies. Journal of Forensic Sciences, 62(4), 986–993. 10.1111/1556-4029.13390

Krebs, C. J. (1999). Ecological Methodology., 2nd edn. Benjamin/Cummings: Menlo Park, CA.

LeBlanc, K., Boudreau, D. R., & Moreau, G. (2021). Small bait traps may not accurately reflect the composition of necrophagous Diptera associated to remains. Insects, 12(3), 261. 10.3390/insects12030261

Losier, C., Boudreau, D. R., LeBlanc, K., Michaud, J.-P., & Moreau, G. (2023). Fall decay deceleration in northern latitudes: Merely a matter of cold? Journal of Medical Entomology, 60(1), 32–39. 10.1093/jme/tjac159

Lutz, L., Amendt, J., & Moreau, G. (2018). Carcass concealment alters assemblages and reproduction of forensically important beetles. Forensic Science International, 291, 124–132. 10.1016/j.forsciint.2018.08.014

Lutz, L., Moreau, G., Czuprynski, S., Bernhardt, V., & Amendt, J. (2019). An empirical comparison of decomposition and fly colonisation of concealed carcasses in the Old and New World. International Journal of Legal Medicine, 133(5), 1593–1602.

Matuszewski, S., Hall, M. J. R., Moreau, G., Schoenly, K. G., Tarone, A. M., & Villet, M. H. (2020). Pigs vs people: The use of pigs as analogues for humans in forensic entomology and taphonomy research. International Journal of Legal Medicine, 134(2), 793–810. 10.1007/s00414-019-02074-5

McAlpine, J. F. (1981). *Manual of Nearctic Diptera*. Volume 1: Monograph: 27 / Agriculture Canada. Canadian Government Publishing Centre.

Mégnin, P. (1887). La faune des tombeaux. Compte Rendu Hebdomadaire des Séances de l’Académie des Sciences, 105, 948–951.

Megyesi, M., Nawrocki, S., & Haskell, N. (2005). Using accumulated degree-days to estimate the postmortem interval from decomposed human remains. Journal of Forensic Sciences, 50(3), 1–9. 10.1520/JFS2004017

Michaud, J.-P., Schoenly, K. G., & Moreau, G. (2012). Sampling Flies or Sampling Flaws? Experimental Design and Inference Strength in Forensic Entomology. Journal of Medical Entomology, 49(1), 1–10. 10.1603/ME10229

Michaud, J.-P., & Moreau, G. (2013). Effect of variable rates of daily sampling of fly larvae on decomposition and carrion insect community assembly: Implications for forensic entomology field study protocols. Journal of Medical Entomology, 50(4), 890–897. 10.1603/ME12200

Michaud, J.-P., Schoenly, K. G., & Moreau, G. (2015). Rewriting ecological succession history: Did carrion ecologists get there first? The Quarterly Review of Biology, 90(1), 45–66. 10.1086/679763

Michaud, J. P., & Moreau, G. (2017). Facilitation may not be an adequate mechanism of community succession on carrion. Oecologia, 183(4), 1143–1153.

Moffatt, C., Simmons, T., & Lynch-Aird, J. (2016). An improved equation for TBS and ADD: establishing a reliable postmortem interval framework for casework and experimental studies. Journal of Forensic Sciences, 61, S201–S207.

Moreau, G., Lutz, L., Boudreau, D. R., Michaud, J. P., & Schoenly, K. G. (2025). Design and analysis of field studies in carrion ecology. In Carrion Ecology, Evolution, and Their Applications (pp. 63–85). CRC Press.

Payne, J. A. (1965). A summer carrion study of the baby pig *Sus Scrofa* Linnaeus. Ecology, 46(5), 592–602. 10.2307/1934999

Pohjoismäki, J. L., Karhunen, P. J., Goebeler, S., Saukko, P., & Sääksjärvi, I. E. (2010). Indoors forensic entomology: colonization of human remains in closed environments by specific species of sarcosaprophagous flies. Forensic Science International, 199, 38– 42.

R Core Team (2025). R: A Language and Environment for Statistical Computing_. R Foundation for Statistical Computing, Vienna, Austria. https://www.R-project.org/

Ratcliffe, B. C. (1972). The Natural History of Necrodes Surinamensis (Fabr.) (Coleoptera: Silphidae). Transactions of the American Entomological Society (1890-), 98(4), 359–410. http://www.jstor.org/stable/25078118

Reibe, S., & Madea, B. (2010). How promptly do blowflies colonise fresh carcasses? A study comparing indoor with outdoor locations. Forensic Science International, 195(1– 3), 52–57. 10.1016/j.forsciint.2009.11.009

Simmons, T., Adlam, R. E., & Moffatt, C. (2010). Debugging decomposition data— Comparative taphonomic studies and the influence of insects and carcass size on decomposition rate. Journal of Forensic Sciences, 55(1), 8–13. 10.1111/j.1556-4029.2009.01206.x

Skidmore, P. (1985). The biology of the muscidae of the world. W. Junk Publishers.

Statistique Canada. (2022). L’homicide au Canada, 2021. Juristat. no 85-002-X. https://www150.statcan.gc.ca/n1/fr/pub/85-002-x/2022001/article/00015-fra.pdf

Syamsa, R. A., Omar, B., Ahmad, F. M. S., Hidayatulfathi, O., & Shahrom, A. W. (2017). Comparative fly species composition on indoor and outdoor forensic cases in Malaysia. Journal of Forensic and Legal Medicine, 45, 41–46.

Tantawi, T. I., El-Kady, E. M., Greenberg, B., & El-Ghaffar, H. A. (1996). Arthropod Succession on Exposed Rabbit Carrion in Alexandria, Egypt. Journal of Medical Entomology, 33(4), 566–580. 10.1093/jmedent/33.4.566

Thümmel, L., Lutz, L., Geissenberger, J., Pittner, S., Heimer, J., & Amendt, J. (2023). Decomposition and insect succession of pig cadavers in tents versus outdoors – A preliminary study. Forensic Science International, 346, 111640. 10.1016/j.forsciint.2023.111640

Turner, M. G., Baker, W. L., Peterson, C. J., & Peet, R. K. (1998). Factors influencing succession: lessons from large, infrequent natural disturbances. Ecosystems, 1(6), 511–523. 10.1007/s100219900047

Valiela, I. (1974). Composition, food webs and population limitation in dung arthropod communities during invasion and succession. American Midland Naturalist, 92(2), 370–385. 10.2307/2424302

Voss, S. C., Forbes, S. L., & Dadour, I. R. (2008). Decomposition and insect succession on cadavers inside a vehicle environment. Forensic Science, Medicine, and Pathology, 4(1), 22–32. 10.1007/s12024-007-0028-z

Weiher, E., & Keddy, P. A. (1995). The assembly of experimental wetland plant communities. Oikos, 73(3), 323–335. 10.2307/3545956

Yan, W., Zhong, Y., Zhu, G., Liu, W., & Shangguan, Z. (2020). Nutrient limitation of litter decomposition with long-term secondary succession: Evidence from controlled laboratory experiments. Journal of Soils and Sediments, 20(4), 1858–1868. 10.1007/s11368-019-02523-z

